# Fyn kinase inhibition using AZD0530 improves recognition memory and reduces depressive-like behaviour in an experimental model of Parkinson’s disease

**DOI:** 10.1101/2021.06.16.448746

**Authors:** Bianca Guglietti, Laura Carr, Benjamin Ellul, Sanam Mustafa, Frances Corrigan, Lyndsey E Collins-Praino

**Author notes:** Address Correspondence To: A/Prof Lyndsey Collins-Praino Helen Mayo North, N212, The University of Adelaide, Adelaide, South Australia 5044 Australia.

## Abstract

Fyn kinase has recently been established as a major upstream regulator of neuroinflammation in PD. This study aimed to determine if inhibition of Fyn kinase could lead to reduced neuroinflammation and improvements in motor and non-motor impairments in an early-stage model of PD. An experimental model of PD was produced using intra-striatal injection (4µl) of the neurotoxin 6-OHDA (5µg/µl). Sprague Dawley rats (n=42) were given either vehicle, 6mg/kg or 12mg/kg of Fyn kinase inhibitor (AZD0530) daily for 32 days via oral gavage and tested on a battery of tasks assessing motor, cognitive and neuropsychiatric outcomes. AZD 0530 administration led to improvement in volitional locomotion and recognition memory, as well as a reduction in depressive-like behaviour. Pathologically, an inflammatory response was observed; however, there were no significant differences in markers of neuroinflammation between treatment groups. Taken together, results indicate a potential therapeutic benefit for use of Fyn kinase inhibition to treat non-motor symptoms of PD, although mechanisms remain to be elucidated.

**HIGHLIGHTS:**

- Fyn kinase has recently been proposed as a major upstream regulator of microglial activation in Parkinson’s disease (PD).
- This study was the first to evaluate the effects of Fyn kinase inhibition in a rodent model of PD.
- Fyn kinase inhibition using the Fyn kinase inhibitor AZD 0530 was capable of improving volitional locomotion and recognition memory and reducing depressive-like behaviour in a rodent model of PD.
- Interestingly, while increases in microglial activation were observed in this rodent model of PD, AZD 0530 did not significantly reduce this activation.
- This suggests that the behavioural improvements associated with Fyn kinase inhibition may occur independently of neuroinflammation and may be attributable to other brain mechanisms, including actions on NMDA or 5-HT_6_ receptors.

## 1. INTRODUCTION

Parkinson’s disease (PD) is the second most common neurodegenerative disease after Alzheimer’s Disease (AD) and the fastest growing neurological disorder(Dorsey et al., 2018; Feigin et al., 2019). Pathologically, PD is characterised by the presence of Lewy Bodies, intracellular aggregates of alpha-synuclein (α-syn) and ubiquitin, and the loss of dopaminergic (DA) neurons in the substantia nigra *pars compacta* (SNpc) (Pollanen et al., 1993). Clinically, the cardinal motor symptoms include muscle rigidity, akinesia, resting tremor, bradykinesia and postural/gait changes (Barnum and Tansey, 2012). PD patients also experience a variety of non-motor symptoms, such as anxiety, depression and cognitive impairment (Barnum and Tansey, 2012). These neuropsychiatric and cognitive impairments significantly affect quality of life and, unfortunately, current treatment strategies are not effective in addressing these symptoms, which are considered amongst the most debilitating (Duncan et al., 2014). Additionally, the most commonly used strategy targeting cognitive impairment, the cholinesterase inhibitors used for AD, may paradoxically worsen motor deficits (Collins-Praino et al., 2011). Accordingly, new therapeutic strategies to address this gap are necessary; however, to date, attempts to develop such treatments have been limited due to an incomplete understanding of the pathogenesis and the molecular mechanisms underpinning cognitive impairment and mood dysfunction in PD.

Neuroinflammation has been proposed as a key factor driving neurodegeneration in PD (Tansey and Goldberg, 2010). McGeer and colleagues first observed the presence of activated microglia in the SN of post-mortem PD patients (McGeer et al. (1988), and, since this observation, many cell culture, animal and port-mortem studies have confirmed the role of microglial-mediated inflammation in PD (Lee et al., 2009; McGeer and McGeer, 2004; Tansey and Goldberg, 2010). Microglia are the first line of CNS immune defence, acting as molecular sensors and surveying the environment where they become ‘activated’ in response to insult or injury, undertaking phagocytosis to maintain homeostasis (Yang et al., 2021). During the early stages of PD, microglia become activated by various signalling pathways in response to stimuli such as protein aggregation and oxidative stress (McGeer and McGeer, 2004). Upon activation, they release ROS, chemokines and pro-inflammatory cytokines, such as TNFα, IL-1β and IFNγ (Yan et al., 2014). Increases in these detrimental cytokines have consistently been reported in the serum and cerebrospinal fluid of PD patients (Mogi et al., 2000). Under physiological conditions, this response is associated with scavenging and healing and terminates; however, if the stimulus persists, chemokines may initiate an adaptive immune response, recruiting more macrophages, leading to an accumulative increase in pro-inflammatory cytokine release (Salvi et al., 2017). Synergistically, pro-inflammatory cytokines bind to their respective receptors on the microglial surface, further accelerating activation and sustaining a chronic inflammatory state (Lee et al., 2002). This chronic inflammatory state is upregulated in PD, with increased microglial density observed in the SN of PD patients (Beach et al., 2007). Overall, this chronic inflammation is detrimental long-term, initiating cell damage and death and exacerbating disease progression in PD (Tansey and Goldberg, 2010). Although the exact causes of initial microglial stimulation and subsequent upregulation in PD are unknown, an upstream element that may be key to understanding these mechanisms is Fyn kinase.

Fyn is an SRC-family kinase (SFK) involved in several key central nervous system (CNS) functions, including myelination, astrocyte migration and SP (Matrone et al., 2020). Fyn also plays a critical role in both peripheral and central inflammatory processes (Nygaard et al., 2014). This has led to speculation Fyn dysregulation may be implicated in the pathophysiology of multiple neurodegenerative diseases, including PD (Angelopoulou et al., 2021; Moore et al., 2002; Panicker et al., 2015; Stuart et al., 2007). In support of this, a seminal study by Panicker and colleagues in both cell culture and animal models of PD, described the critical role of Fyn in regulating the microglial-mediated inflammatory response through nuclear translocation of the p65 component of NFκß into the nucleus and subsequent up-regulation of pro-inflammatory cytokine production (Panicker et al., 2015). Fyn has also been implicated in the signalling pathway linking αsyn oligomers to neurodegenerative disease processes in PD via its binding to the cellular prior protein receptor (PrPc), leading to Fyn activation and subsequent upregulation of NMDA induced excitotoxicity (Ferreira et al., 2017). Recently, a GWAS study identified the FYN gene as a novel PD risk locus (Nalls et al., 2019), with the Fyn/PKCδ signalling pathway known to contribute to oxidative-stress induced death of dopaminergic neurons (Kaul et al., 2005; Saminathan, 2011). Indeed, in multiple mouse models of PD (LPS, MPTP and 6-OHDA), a greater attenuation of the neuroinflammatory response was seen in Fyn knockout mice compared to wild-type (Panicker et al., 2015), further supporting the role of Fyn as a major upstream regulator of the neuroinflammatory processes in PD. Inhibition of Fyn may therefore represent a novel target for therapeutic intervention. To date, no studies have investigated the benefits of Fyn kinase inhibition in a preclinical model of PD.

AZD0530 (Saracatinib) is an experimental drug which acts as an inhibitor of Src Family Kinases (SFKs), with high specificity for Fyn (Hennequin et al., 2006). Initially developed for the treatment of solid tumours, the drug demonstrated success in animal models; however, was unable to translate to humans, potentially due to the high threshold required for kinase inhibition to modify tumour progression (>98%) (Nygaard et al., 2014). AZD0530 has since been Arepurposed for its potential therapeutic use in neurodegenerative diseases, which require a much lower level of kinase inhibition to modify the inflammatory pathway (Nygaard et al., 2014). Recently, AZD0530 was found to be efficacious in a transgenic mouse model of AD, fully rescuing spatial memory deficits and synaptic depletion and dampening microglial activation (Kaufman et al., 2015). The safety, tolerability and availability of AZD0530 has also been assessed in humans with AD and was found to be safe and well tolerated across doses (Nygaard et al., 2015). The drug has not yet been assessed in PD; however, given its utility in other neurodegenerative diseases, AZD0530 may represent a novel candidate for therapeutic use in the disease. Here, we investigate whether AZD0530 can be repurposed for use in PD by evaluating its effect both on parkinsonian motor impairment, as well as non-motor symptoms of the disease, including cognitive and neuropsychiatric dysfunction, using the striatal 6-OHDA model of PD.

## 2. MATERIALS AND METHODS

### 2.1 Animals

Adult male Sprague-Dawley rats weighing between 280-360g (*n=42*) were used in this study. All experimental procedures were approved by the of the University of Adelaide Animal Ethics Committee (M-2015-241a) and conducted according to the Australian National Health and Medical Research Council code of practice for the care and use of animals for scientific purposes (8th edition, 2013). Animals were housed under conventional laboratory conditions with a 12-hour light/dark cycle and access to standard rodent chow and water *ad libitum*. All experiments were conducted in the AM during the light phase of the light/dark cycle.

To represent PD, a unilateral striatal 6-hydroxydopamine (6-OHDA) model was adopted (Schwarting and Huston, 1996). 6-OHDA has a high affinity for dopamine transporters (DAT), which transport the toxin into the cell, causing mitochondrial dysfunction and oxidative stress and leading to cell death (Ungerstedt et al., 1974). The intra-striatal approach causes 30-40% loss of striatal DA content, representing early-stage PD (Yuan et al., 2005) and is associated with significant increases in neuroinflammatory markers (Ramsey and Tansey, 2014).

### 2.2 Stereotactic Surgery

Animals were anaesthetised via inhalation of 5% isoflurane under normoxic conditions (0.6 L/min 02, 1.5 L/min N_2_) and mounted securely on a stereotactic frame in a flat skull position with the incisor bar 3.0 mm below horizontal zero. Anaesthesia was maintained with 2.5% inhalational isoflurane via nose cone. Lignocaine (0.25 ml) was administered prior to a midline incision and soft tissue subsequently retracted to expose bregma. Animals were randomly assigned to receive either left or right striatal lesions. Based on methods previously described (Lee et al., 1996; Thornton and Vink, 2012), a 1mm burr hole was made at two separate locations over the striatum using the following standard stereotactic coordinates: (1) AP: 0.5 mm, ML: +/- 2.5 mm, DV: −5.0 mm and (2) AP: −0.5 mm, ML: +/- 4.2, DV: −5.0 mm. Two μl of 6-OHDA (Sigma) (5 μg/μl) was injected at a rate of 1 μl/min into each coordinate (4 μl in total) using a 25 μL Hamilton syringe, which was left in place for 2 minutes to allow for diffusion. Sham animals received equivalent injection of 0.9% sodium chloride (saline). Following injections, anaesthesia was ceased, the incision was closed using 9 mm surgical autoclips and betadine was applied to wound site. Animals were placed on a heatpad to thermostatically maintain body temperature both during and post-surgery and monitored for normal recovery.

### 2.3 Experimental Design and Drug Administration

Animals were randomly assigned into 4 experimental groups *(n=10-11)* and treated with vehicle control (sham + veh; 6-OHDA + veh), 6 mg/kg/day (6-OHDA + 6 mg/kg) or 12 mg/kg/day (6-OHDA + 12 mg/kg) of AZD0530 daily for 33 days via oral gavage. All treatments were administered in a blinded fashion, with allocation of the doses prepared daily by a blinded independent researcher. The vehicle was 0.5% wt/vol hydroxypropyl-methylcellulose (HPMC)/0.1% wt/vol polysorbate 80. All animals were dosed in the AM and, on days of functional testing, were treated 1 hour prior to commencement. Treatment doses were chosen based upon efficacy in pilot work and previous studies in preclinical models of AD (Kaufman et al., 2015).

### 2.4 Functional Testing

Animals were tested in the AM for all functional assessments and recorded using the ANY-Maze video tracking system (Stoelting Co. V.4.99m) under the following regime (Figure 1). The experimenter was blinded to experimental groups throughout the study duration.

**Figure 1.**
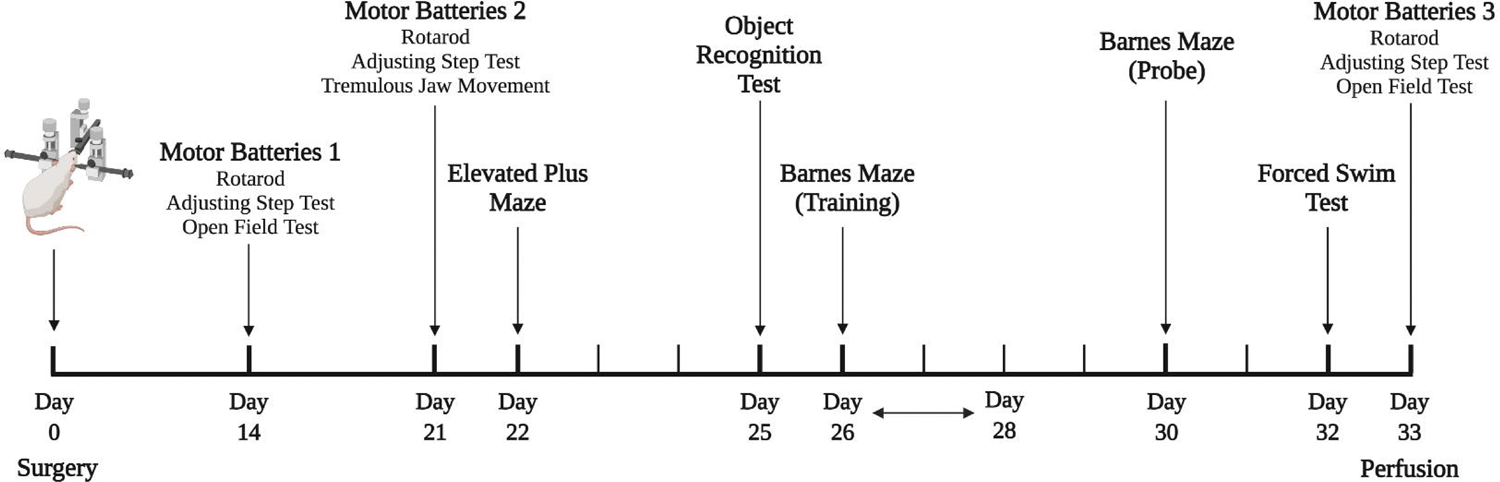
Overview of Experimental Design Timeline. Figure created in BioRender.com (2021).

#### 2.4.1 Rotarod

Rotarod is a common test assessing co-ordination, balance and general locomotion (Deacon, 2013). The device comprises an elevated circular set of metal horizontal bars which rotate on a longitudinal axis between 0-30 revolutions per minute (RPM). Animals were placed on the stationary rotarod for 10 seconds before speed was increased by 3 RPM every 10 seconds until a maximum of 30 RPM. Animals maintained the speed for a further 20 seconds before the speed was decelerated and the animal removed from the test. The test concluded either when the animal fell, gripped without walking for 2 complete revolutions or reached the full 120 seconds and latency was recorded.

#### 2.4.2 Adjusting Step Test

The adjusting step test (AST) is a test assessing forelimb akinesia and gait impairments (Olsson et al., 1995). Animals were held so their hind limbs and the ipsilateral forepaw were unable to touch the testing surface. With only the contralateral forepaw touching, animals were moved laterally across a distance of 45 cm. The number of adjusting steps were recorded for contralateral forehand and backhand by slowly moving the animal across the testing surface 3 times in either direction. The average number of adjusting steps taken across the trials was calculated.

#### 2.4.3 Open Field Test

The Open Field Test (OFT) incorporates assessment of both motor and anxiety-like behaviour (Leite-Almeida et al., 2009). Animals were placed in the centre of a large square box (95 x 95 cm) with walls at a height of 44.5 cm for 5 minutes. The total distance travelled whilst in this space was measured to assess locomotion and time spent in inner zone measured to assess anxiety.

#### 2.4.4 Tremulous Jaw Movement

Tremulous jaw movements (TJM) in rodents occur in the same local frequency as parkinsonian tremor (3-7Hz) (Collins-Praino et al., 2011). The TJM test is thus a validated method of assessing parkinsonian tremor in the 6-OHDA rodent model (Jicha and Salamone, 1991). To assess this, animals were placed in a clear ventilated container (29cm X 20cm X 16cm), allowing continuous visualisation of the lower jaw from all angles. An initial habituation period of 5 minutes was undertaken, followed by two testing phases of 5 minutes. During testing, animals were continuously observed, and any non-directed movement of the lower jaw was recorded via a mechanical hand counter by a trained and blinded observer. An average across both testing phases was taken to indicate tremor severity.

#### 2.4.5 Elevated Plus Maze

The elevated plus maze (EPM) is a widely used tool to assess anxiety-like behaviour in rodents (Pego et al., 2008). Animals were placed in the junction of a cross-shaped elevated maze (50cm in height) with two open (50 cm length) and two closed arms (40cm high × 50 cm length) for 5 minutes. Time spent in the closed arms versus open arms was recorded, with increased time spent in the closed arms representing anxiety-like behaviour.

#### 2.4.6 Novel Object Recognition

The novel object recognition test (ORT) is a common method to investigate recognition memory in rodents (Ennaceur and Delacour, 1988). For this study, an abridged version of the standard ORT test was used. Here, animals were placed in an empty arena (100cm × 45cm) for 5 minutes to habituate. Thirty minutes later, they were placed back in for 5 minutes with 2 objects. Time spent inspecting each object was measured. Inspection was operationally defined as approaching, looking at and sniffing the object from a distance of less than or equal to 2cm. In order to assess long-term reference memory, 24 hours later, animals were placed back in the arena with 1 old object and one ‘novel’ object. Time spent inspecting both objects was measured. To determine novel object preference, time spent inspecting the novel object compared to the old object was converted to a preference by dividing time spent with novel object by overall time inspecting both objects and converted to a percentage ([time_novel_/ time_familiar_ + time_novel_] *100). A score of greater than 50% indicates a preference for the novel object.

#### 2.4.7 Barnes Maze

Barnes Maze is a well-known measure of spatial learning and memory, as well as cognitive flexibility (Sunyer et al., 2007). The apparatus consists of a circular table (1.2 m in diameter) with 18 circular holes equally spaced around the perimeter edge. One of the holes was pre-designated as an escape hole, with a black escape box placed immediately below the hole. The Barnes maze task was performed over the course of five days: three days of acquisition trials (Days 1-3); a rest day, during which there was no interaction with the animals (Day 4) and a probe day, in which the previous location of the escape box was changed to a new location (Day 5). During acquisition days, animals underwent two trials (spaced 15 minutes apart). On all trials, animals were placed in the centre of the Barnes maze with a bright light shone overhead as an aversive stimulus, motivating animals to find and enter the escape box. The latency for animals to find and enter the escape box was recorded. On the probe day, the location of the escape box was changed to a new hole. Two trials were then conducted, one hour apart. In the first trial, the time taken to reach the old location of the escape box was recorded as a measure of long-term spatial reference memory. On both probe day trials, the latency to find and enter the new location of the escape box was recorded as a measure of cognitive flexibility.

#### 2.4.8 Forced Swim Test

The forced swim test (FST) is a method for assessing depressive-like behaviour in rats (Slattery and Cryan, 2012). A clear cylindrical vase was half filled with 25 °C water, adjusted for the animal’s length so that the hind legs did not touch the bottom of the cylinder. Rats were placed in the tank for 5 minutes and time spent swimming and immobile were measured. Increased time spent immobile indicates behavioural despair, a measure of depressive-like behaviour.

### 2.5 Tissue collection and Processing

At day 33, rats were anaesthetised with 5% isoflurane before being randomly assigned for either immunohistochemical (n=21) or molecular analysis (n=21).

Animals that were to be used for immunohistochemical analysis were transcardially perfused with 0.2mL heparin + 10% formalin. Brains were removed and post-fixed in 10% formalin for one week, then blocked into 2 mm coronal sections and embedded in paraffin-wax. Five μm coronal sections of the striatum beginning +0.5mm from Bregma were prepared from paraffin embedded tissue. Tissue was mounted on silane coated slides and allowed to dry at 37 °C overnight.

Animals that were to be used for molecular analysis were transcardially perfused with 0.9% saline and the brain extracted whole (n=4-6 per group). Brains were snap-frozen in liquid nitrogen, then stored at −80° C. Prior to analysis, brains were placed on dry ice and tissue samples from the striatum were dissected. Tissue was suspended in buffer solution (20u Tris-HCl pH 7.5, 2u u mM 2-mercaptoethanol) with protease inhibitor cocktail (Sigma), 10 μg/mL aprotinin, leupeptin, pepstatin A and 10u PMSF. Samples were then homogenised and sonicated in 3 × 10 s bursts using a sonicator probe. Following this, homogenised samples were centrifuged for 15 minutes at 14,000 rpm at 4° C, before supernatant was collected. Protein concentrations were estimated with Thermo Pierce BCA Protein Assay Kit (ThermoScientific) at 540 nm absorbance.

### 2.6 Immunohistochemical Analysis

Immunohistochemical analysis was performed using standard protocols. Briefly, following deparaffinization in xylene, tissue mounted slides were rehydrated in ethanol and endogenous perioxidases blocked using methanol with 0.5% hydrogen peroxide. Slides were then washed twice with phosphate buffered saline (PBS). Antigen retrieval was performed (citrate), following which sections were blocked in 3% normal horse serum (NHS) for 30min before overnight incubation with Tyrosine hydroxylase (TH) (Abcam, ab112, 1:750) or IBA-1 (Wako, 019-19741, 1:800). On the following day, slides were again washed twice in PBS and incubated with secondary antibody (Vector Goat Anti-Rabbit IgG, 1:250 NHS) for 30 minutes, followed by two more washes with PBS and incubation with streptavidin peroxidase conjugate (Vector SPC,1:1000 NHS) for 1 hour. Following a final wash with PBS, the chromogen 3,3-Diaminobenzidine tetrahydrochloride (Vector DAB; 1:50) was applied for 7 minutes. Slides were counterstained with haematoxylin, dehydrated in ethanol and cleared with xylene, prior to being coverslipped.

Following staining, slides were scanned using a Nanozoomer slide scanner (Hamamatsu, Shizouka, Japan) and the associated software (NDPview, version 2) was used to view images. TH in the striatum was assessed in ImageJ using colour deconvolution to remove haematoxylin and converted to binary. The striatum was isolated and the % positive area was then calculated for each hemisphere by 2 assessors blinded to experimental conditions. The % difference between ipsilateral and contralateral hemispheres was then calculated for 2 slides per sample to reveal the average % loss (as below). An average of the 2 assessors average % loss was calculated for final analysis.

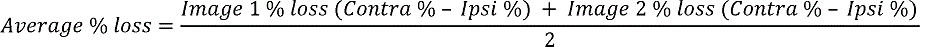

IBA-1 was assessed by counting the reactive and immunopositive cells per mm^2^ in the striatum. Cells were counted by extracting images from the ipsilateral striatum (4 × boxes at 1.04mm^2^ each). These were processed via colour deconvolution, converted to binary and a threshold was applied to demarcate IBA-1+ cells. An automatic cell counter was used and this count was averaged over 2 sections and divided by the total area to generate the number of cells per mm^2^. This was then repeated and compared to the contralateral hemisphere and the difference between the two was calculated. Microglial activation was assessed manually by randomly selecting 6 boxes (3 × 2), across the striatum, each with an area of 0.208mm^2^. Morphological characteristics were assessed, where activated microglia were characterised based on dark nuclear staining, round cell body and short/absent processes. The experimenter was blinded to the experimental group during microglial assessment.

### 2.7 Western Blot Analysis

Western blot analysis was performed using a standard milk method. In brief, 4 × Bolt LDS sample buffer and 10× Bolt reducing agent and dH_2_0 were added to supernatant (30 μ as per the manufacturer’s instructions, heated at 70° C for 10 minutes and then vortexed. Gel electrophoresis was performed using Bolt 4-12% Bis-Tris gels (Life Technologies) with 40μ of sample loaded per well. Gels were run at 120 V for 120 minutes. Following this, gels were transferred to the PVDF membrane using the iBlot 2 Dry Blotting System (Life Technologies). Membranes were washed in 1 × tris-buffered saline (3 washes × 5 min), stained with Ponceau S red solution (Fluka Analytical) for 5 minutes to allow for protein visualisation (in order to ensure equal protein loading between lanes), and then washed with dH_2_0 until removal of Ponceau had been achieved.

Membranes were blocked for 2 hours (5% w/v dried skim milk in TBS-T) before overnight incubation at 4° C (2% w/v dried skim milk in TBS-T and primary antibody). Primary antibodies were used at individually optimised concentrations: anti-GFAP (1:40,000, ab7260, Abcam) and the housekeeper chicken anti-GAPDH (1:4000, ab83956, Abcam). Following overnight incubation, membranes were washed in TBS-T (3 × 5 min) and incubated in 2% milk and secondary antibodies, according to the species the primary was raised in (donkey anti-rabbit and donkey anti-chicken, IRDye 800CW; LI-COR, Inc., 1:10000) and incubated for 2 hours at room temperature. Following incubation, each membrane was washed in TBS-T (3 × 5 min) and visualised using an Odyssey Infrared Imaging System (model 9120; software version 3.0.21) (LI-COR, Inc.) at a resolution of 169uμm. Semi-quantitative analysis of band signals was performed using ImageJ version 1.49 and Image Studio Lite version 5.2. Normalisation of blots were performed using a control sample (sham) across the same protein of interest. Relative density of each sample was calculated based on the adjusted density for each blot as described:

Adjusted density = *signal of sample/housekeeper* ***/*** *signal of control protein / housekeeper*

Relative density = *adjusted density of protein* ***/*** *adjusted density of housekeeper*

### 2.8 Statistical Analysis

Data was analysed using Prism software (GraphPad v.7). For some behavioural tests, animals were excluded if they failed to perform the task (n=3 Rotarod, n=2 TJM, n=1 EPM, n=2 ORT). To compare differences between time and treatment groups on rotarod, AST, OFT and Barnes Maze, a repeated measures two-way Analysis of Variance (ANOVA) was used with time as a within subjects factor and group as a between subjects factor. Post hoc testing was conducted according to Bonferroni’s method. Where a significant main effect of time was noted, a one-way ANOVA was run at each timepoint to further probe effects. For all other measures, in order to compare differences between treatment groups, data were analysed using one-way Analysis of Variance (ANOVA) with Tukey’s multiple comparison post-hoc test. All values are displayed as Mean ± SEM, with significance level set at p<0.05.

## 3. RESULTS

### 3.1 Impairments in volitional locomotion and AST were progressive in nature, but no improvements in motor function were observed with AZD0530 treatment

General locomotor activity was measured as time on the rotarod at days 14, 21 and 33 post-surgery (Figure 2A). A significant main effect of time was observed (F_2,68_=28.32, p<0.0001), with all groups improving from day 14 to day 33. No main effect of treatment was found (F_3,34_=0.917, p=0.443), nor was there a significant interaction between testing day and treatment group (F_6,68_=1.038, p=0.409).

**Figure 2.**
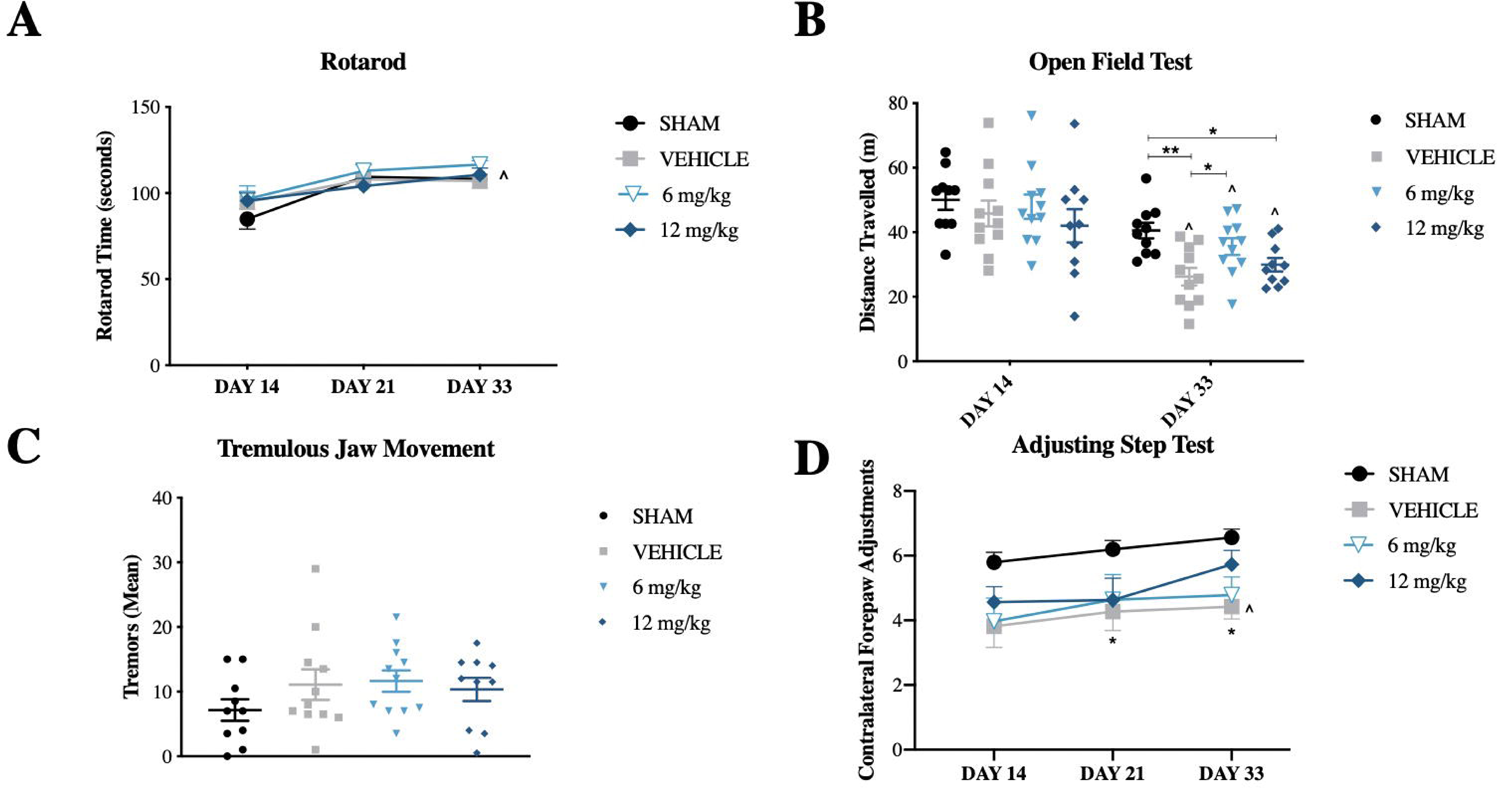
Motor outcomes. (A) Motor function as measured by time on Rotarod (s) at days 14, 21 and 33 post-surgery. (B) Locomotion as measured by distance travelled (m) on Open Field Test at day 14 and 32. (C) Motor impairments as measured by mean number of tremors in Tremulous Jaw Movement. (D) Motor impairments as measured by Contralateral Forepaw adjustments on Adjusting Step Test at days 14, 21 and 32. Graphs represent the Mean± SEM (n=9-11 per group). *p<0.05, **p<0.01 compared to sham (unless indicated otherwise), ^ denotes significance over time.

Volitional locomotion was also assessed as total distance travelled on day 14 and 33 in the OFT (Figure 2B). There was a significant main effect of time (F_1, 38_= 40.77, p<0.0001) with vehicle-(p<0.0001), 6 mg/kg-(p<0.05) and 12 mg/kg-(p=<0.05) treated animals, but not sham animals, travelling significantly less over time. A main effect of treatment was also observed (F_3, 38_= 2.93, p<0.05); however, no significant interaction between time and treatment group was seen (F _3,38_=1.104, p=0.360). Post hoc analysis of treatment effect in the overall model revealed no significant differences between groups (p > 0.05), so, to probe further, a one-way ANOVA was conducted for each individual timepoint. At day 14, no significant differences between shams (50.01±3.07), vehicle-treated (45.82±4.0), 6 mg/kg (47.92±3.79) or 12 mg/kg (42.01±5.18) treated animals were observed (F_3,38_=0.68, p=0.57). At day 33, however, there was a significant difference between sham (40.55±2.43) animals and both vehicle-(26.22±2.72) and 12 mg/kg-treated animals (29.96±2.09) (p<0.001, p<0.05 respectively), indicating an injury effect that was not improved with 12 mg/kg of AZD0530. There was also a significant difference between vehicle and 6 mg/kg (35.57±2.60) treated animals (p<0.05); however, 6 mg/kg treated animals actually travelled significantly more, indicating a potential improvement in locomotion (F_3,38_=6.34, p<0.001).

Tremor was assessed as the mean number of tremors between 2 trials in the Tremulous Jaw Movement (TJM) test on day 21 (Figure 2C). There were no significant differences observed between sham, 6-OHDA vehicle or treatment groups (F_3,38_=1.173, p=0.368).

Akinesia was assessed via the AST (Figure 2D). Mean contralateral forepaw adjustments at day 14, 21 and 33 were analysed. There was a significant main effect of time (F_2,76_=7.8, p<0.001), with post-hoc analysis revealing that, of the 6-OHDA lesioned animals, only 12 mg/kg treated animals demonstrated significantly reduced akinesia over time, with an increased frequency of forepaw adjustments from day 14 (4.56±1.52) to day 33 (5.73±1.377) (p<0.01). A main effect of treatment was also observed (F_3,38_=3.26, p<0.05), with vehicle-treated animals demonstrating akinesia compared to shams at both day 21 (p<0.05) and day 33 (p<0.05), but not day 14 (p > 0.05), indicating a delayed injury effect; however, there was no significant difference between vehicle and any treatment groups (p>0.05). No significant interaction between time and treatment group was observed (F_6,76_=0.58, p=0.75).

### 3.2 Intrastriatal injection of 6-OHDA led to depressive-like behaviour, which was reduced by AZD0530 treatment

Anxiety-like behaviour was assessed using both the EPM and OFT. There was no significant difference in anxiety-like behaviour on the EPM (Figure 3A) between any of the groups on day 22 (F_3,37_= 1.14, p=0.346), indicating no injury or treatment effect. Anxiety-like behaviour, as measured by time in centre of the OFT (Figure 3B) on days 14 (shams 14.56±8.62, vehicle-treated 15.54±8.35, 6 mg/kg 19.72±19.70 or 12 mg/kg 17.15±9.33) and 33 (shams (15.6±8.75), vehicle-treated (8.24±8.87), 6 mg/kg (13.75±10.04) or 12 mg/kg (8.86±4.80)), revealed a significant main effect of time (F_1,38_=6.486, p<0.05); however, there were no significant differences observed upon post-hoc analysis. There was no significant main effect of treatment (F_3,38_=0.70, p=0.558), nor was there a significant interaction between time point and treatment group (F_3, 38_=1.06, p=0.379).

**Figure 3.**
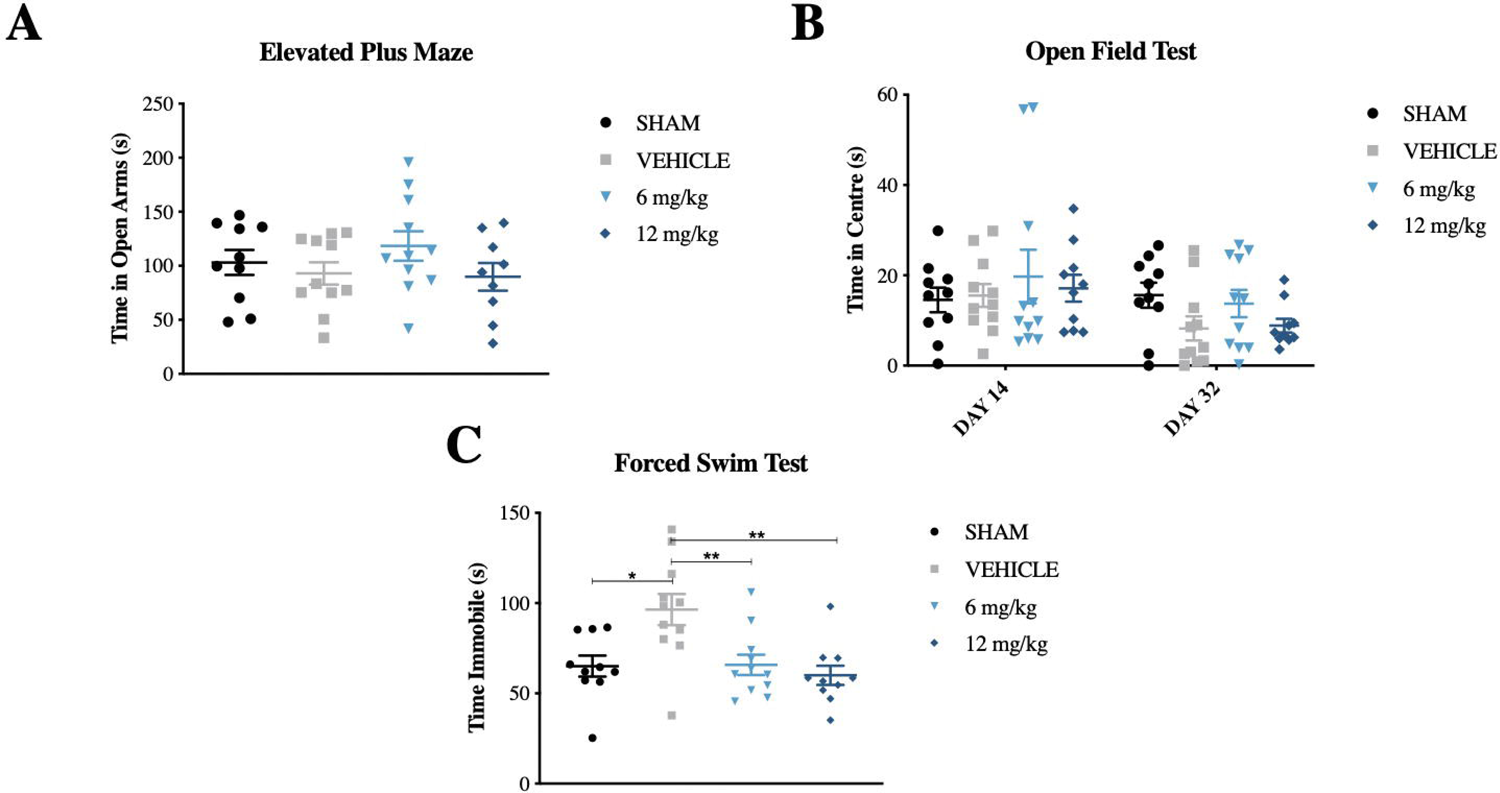
Neuropsychiatric outcomes. **(A)** Anxiety-like behavior as measured by time spent in open arms of Elevated Plus Maze. (B) Anxiety-like behavior as measured by time in centre of Open Field Test at days 14 and 32. (C) Depressive-like behavior as measured by Forced Swim Test. Graphs represent the Mean±SEM (n=10-11 per group). *p<0.05, **p<0.01 compared between injury groups.

Depressive-like behaviour was assessed based on immobility time in the FST (Figure 3C). 6-OHDA vehicle-treated animals spent significantly more time immobile compared to shams (96.41±8.60 vs 65.09±5.81, F_3,38_ =6.570, p<0.01). Vehicles also spent significantly more time immobile than either 6mg/kg (65.81±5.58, p<0.01) or 12 mg/kg (60.02±5.309, p<0.01) treated animals. There was no significant difference between shams and either 6 mg/kg or 12 mg/kg-treated animals (p> 0.05). Together, results indicate a potential reduction in depressive-like behaviour back to sham equivalent following treatment.

### 3.3 AZD0530 administration improved recognition memory as measured by preference for the novel object

Recognition memory was assessed via novel object preference in the ORT (Figure 5.4A). A significant main effect of time was observed (F_1,36_=32.39, p<0.0001), with sham (p<0.05), 6 mg/kg (p<0.05) and 12 mg/kg (p<0.001) treated animals showing an increased preference for the novel object in trial 2, an effect not seen in 6-OHDA vehicle animals. There was also a significant main effect of treatment (F_3,36_=3.133, p<0.05). No significant differences were seen between any of the treatment groups during trial 1 (sham: 47.69±2.81, vehicle: 47.48±3.64, 6 mg/kg: 50.19±5.54 and 12 mg/kg: 48.23±4.916). In trial 2, however, shams had a higher preference index (69.60±5.82) compared to vehicle-treated 6-OHDA animals (52.70±4.633, p<0.05), indicating an injury effect. A treatment effect was also observed, with both 6 mg/kg-treated (70.90±3.21, p<0.05) and 12 mg/kg-treated (75.16±3.90, p<0.01) animals exhibiting increased preference for the novel object compared to vehicle-treated animals.

The BM was used to assess various elements of visuo-spatial learning and memory. Learning acquisition, as assessed during the BM training phase (Figure 4B), demonstrated a significant main effect of time (F_2,76_=18.68, p<0.0001), with vehicle-(p<0.0001), 6 mg/kg-(p<0.001) and 12 mg/kg-treated animals (p<0.05) all improving from day 1 to day 3, demonstrating learning of the task. Shams did not demonstrate significant improvement from day 1 to 3 (p=0.659), likely due to their maximal learning of the task on day 1. There was no significant main effect of treatment (F_3,38_=1.398, p=0.258) and no significant interaction between day and treatment group (F_6,76_ =1.518, p=0.184). Similarly, spatial reference memory, as observed in the latency to old escape box on trial 1 of probe day, revealed no significant differences (F _3,38_= 0.626, p<0.603) between groups (Figure 4C).

**Figure 4.**
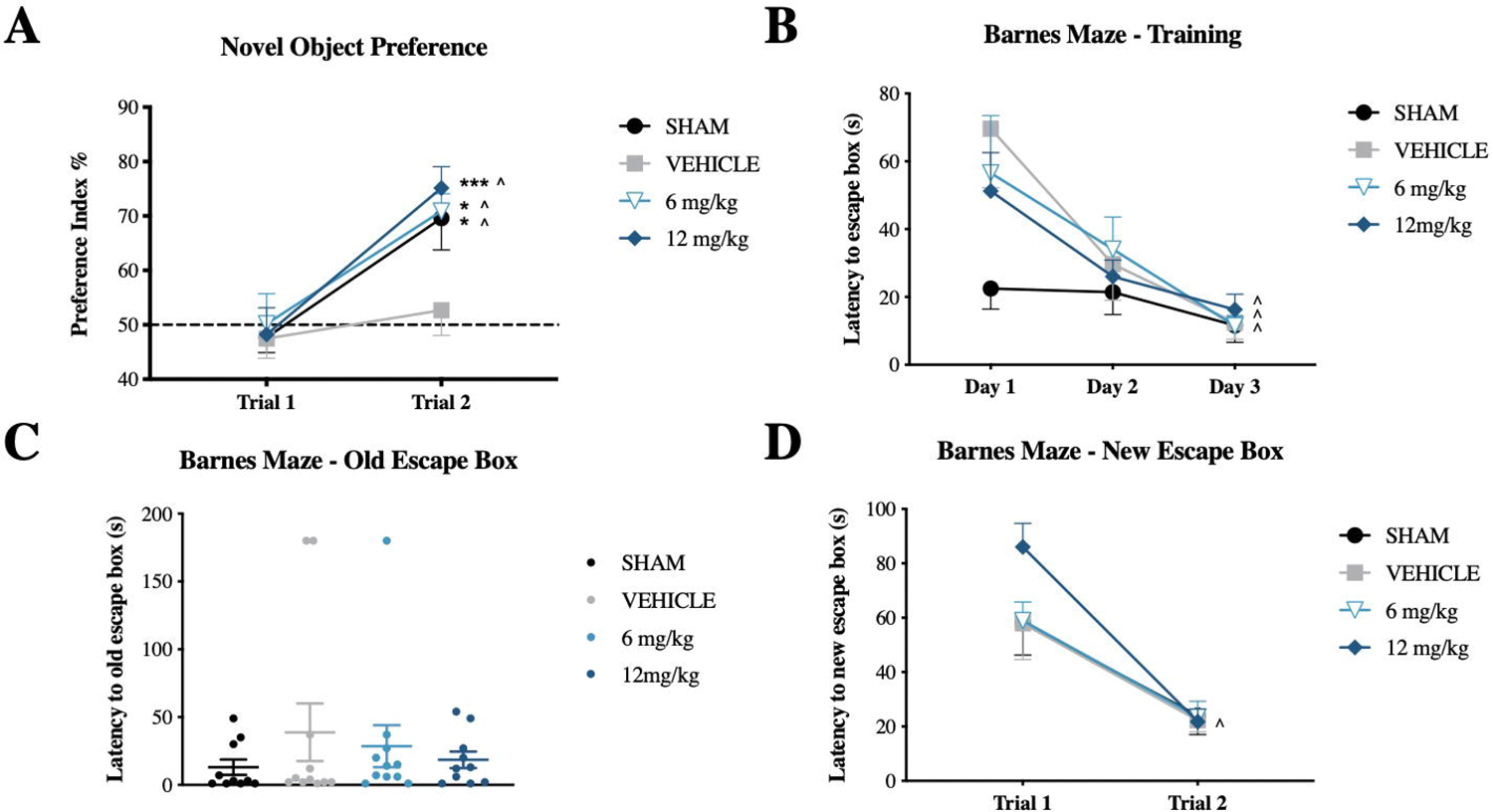
Cognitive outcomes. (A) Novel object preference as assessed by preference index (%) for object B on the Object Recognition Test (B) Visuo-spatial learning as assessed by latency to escape box on training days 1, 2 and 3 on Barnes Maze. (C) Spatial reference memory as assessed by latency to old escape box. (D) Cognitive flexibility measured by latency to new escape box on probe day. Graphs represent Mean± SEM (n=10-11 per group). *p<0.05, ***p<0.001 compared to vehicle, ^ denotes significance over time.

Cognitive flexibility was examined via the ability to learn a new location for the escape hole over two trials on probe day (Figure 5D). There was a significant main effect of time (F _1,38_=52.10, p<0.0001), with all groups demonstrating improved latency to the escape hole from trial 1 to trial 2 (p<0.05). There was no significant main effect of treatment (F_3,38_=1.180, p=0.33) nor interaction between trial and treatment group (F_3,38_=1.50, p=0.23).

**Figure 5.**
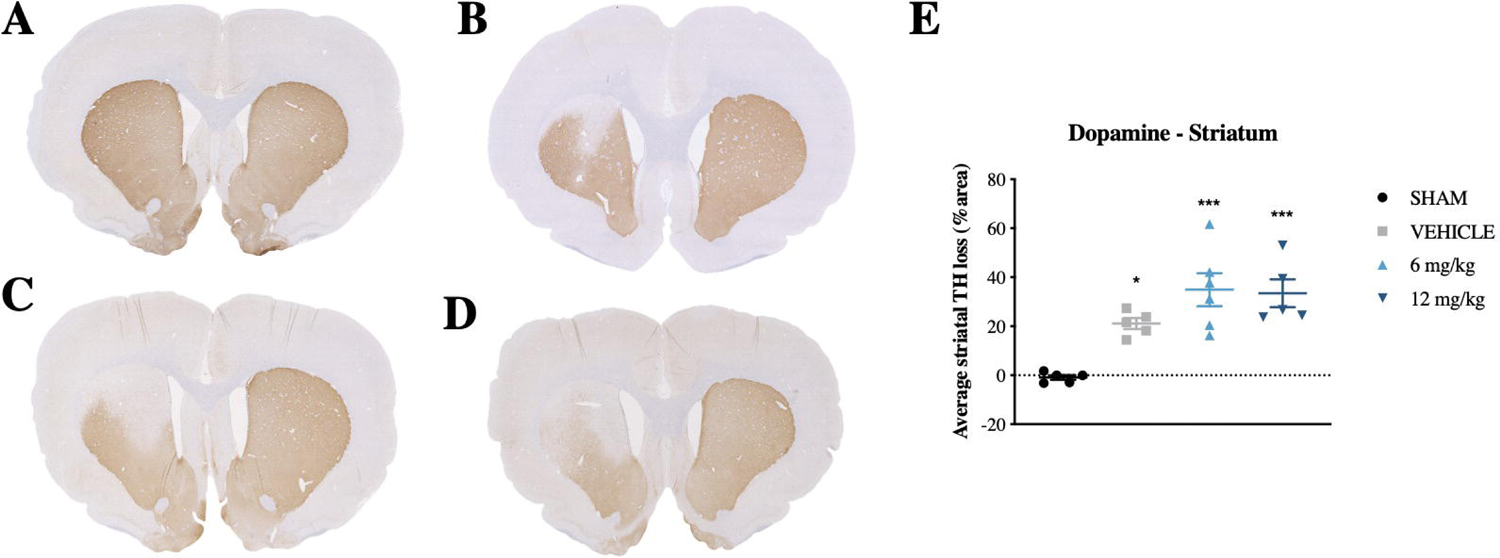
Representative images of TH staining within the striatum (A-D) of Sham (A), Vehicle (B), 6mg/kg (C) and 12mg/kg (D), as well as respective graph (E). Graphs represent Mean± SEM (n=5-6 per group). *p<0.05, ***p<0.001 compared to Sham.

### 3.4 AZD0530 did not rescue dopaminergic integrity in the striatum

Dopaminergic integrity was assessed via % area of TH +ve staining in the ipsilateral relative to contralateral striatum following 6-OHDA injection (Figure 5). Dopaminergic loss in sham animals was unremarkable. A significant loss of dopaminergic integrity was observed (F_3,17_ = 11.51, p<0.001) in vehicle-(21.1±2.24%), 6 mg/kg-(34.94±6.71%) and 12 mg/kg-treated (33.46±5.65%) animals compared to sham (−0.92±0.95%). There were no significant differences between vehicle and either 6 mg/kg or 12 mg/kg treated animals (p> 0.05), indicating no prevention of dopaminergic loss with treatment.

### 3.5 No alterations to measures of inflammation were observed with AZD0530 administration

The neuroinflammatory response was assessed both by average number of IBA1 +ve cells (Figure 5.6) and by activation state in the striatum (Figure 6). There was a significant main effect of group on total microglial population in the striatum (F_3,17_=6.187, p=0.01, with vehicle-(97.14±6.70, p<0.05), 6 mg/kg-(106.9±8.02, p<0.01) and 12 mg/kg-treated animals (108.1±16.16, p<0.01) all displaying a higher average number of microglia compared to shams (49.78±11.47) (Figure 6E). There was no significant difference in microglial numbers between vehicle and either 6 mg/kg- or 12 mg/kg-treated animals (p>0.05), indicating that AZD0530 treatment did not reduce total microglial population.

**Figure 6.**
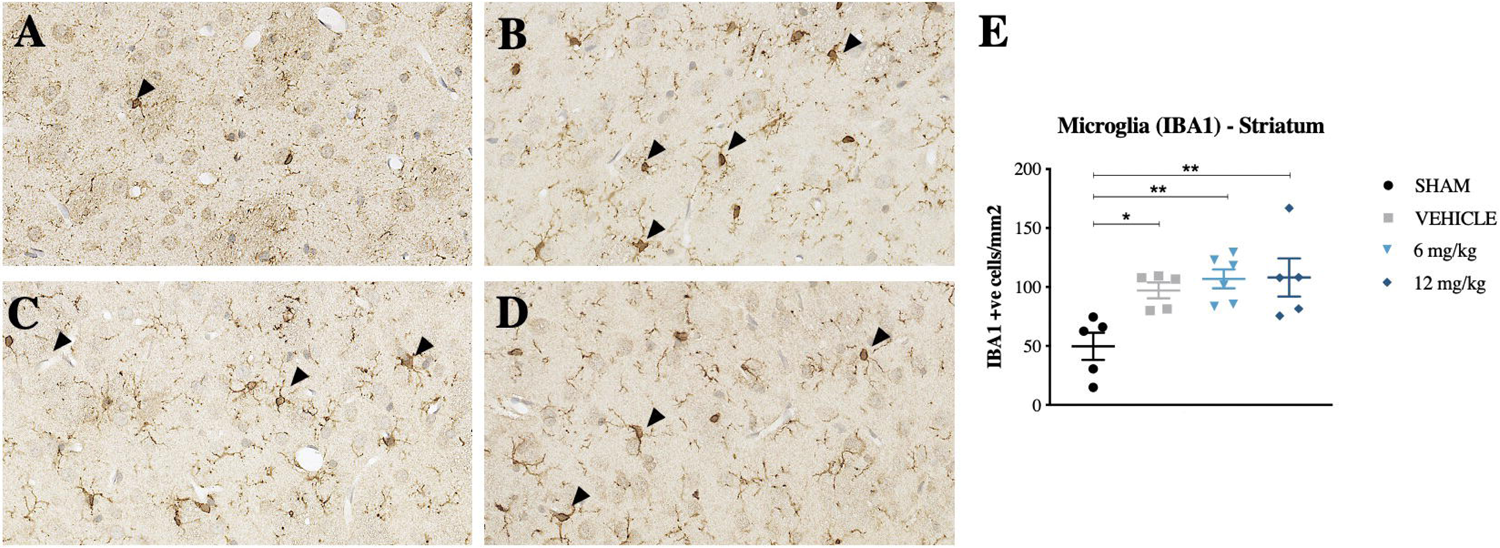
Representative images of IBA1 staining within the striatum (A-D) of Sham (A), Vehicle (B), 6mg/kg (C) and 12mg/kg (D), as well as respective graph (E). Graphs represent Mean± SEM (n=5-6 per group). *p=<0.05, **p=<0.01 compared to Sham.

Activation state was characterised by analysing the cellular morphology of IBA1 +ve cells (Figure 7D). There was a significant main effect of group on activation status of the microglia (F_3,16_=3.638, p=0.04. Specifically, there was a significant increase in the number of activated microglia observed in vehicle-treated (51.41±6.51, p<0.05) animals when compared to shams (20.13±4.00). Interestingly, microglial activation was not increased in either the 6 mg/kg-(46.07±5.93, or 12 mg/kg-(39.10±8.66) treated animals compared to shams (p>0.05). However, the difference in microglial activation between vehicle and either 6 mg/kg- or 12 mg/kg-treated animals was not significant (p>0.05).

**Figure 7.**
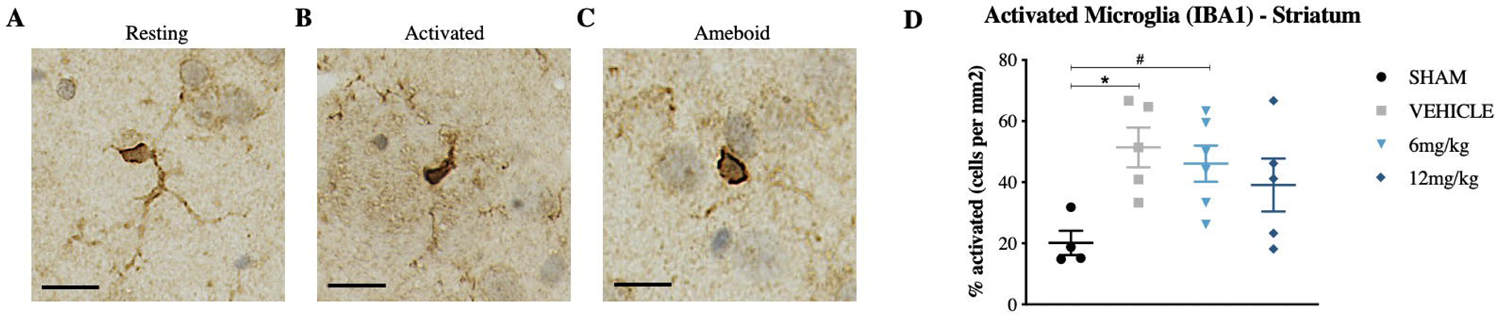
Representative images of morphological appearance of IBA1 staining of (A) resting, (B) activated and (C) ameboid. Scale bar represents 25μm. Respective graph represents activated microglia (B + C relative to total). Graphs represent Mean± SEM (n=5-6 per group). *p=<0.05, #p=<0.01 compared to Sham.

Astrocytic response was assessed via GFAP levels in western blot (Figure 8). No significant main effect of group was found (F _3,15_=0.865, p=0.481).

**Figure 8.**
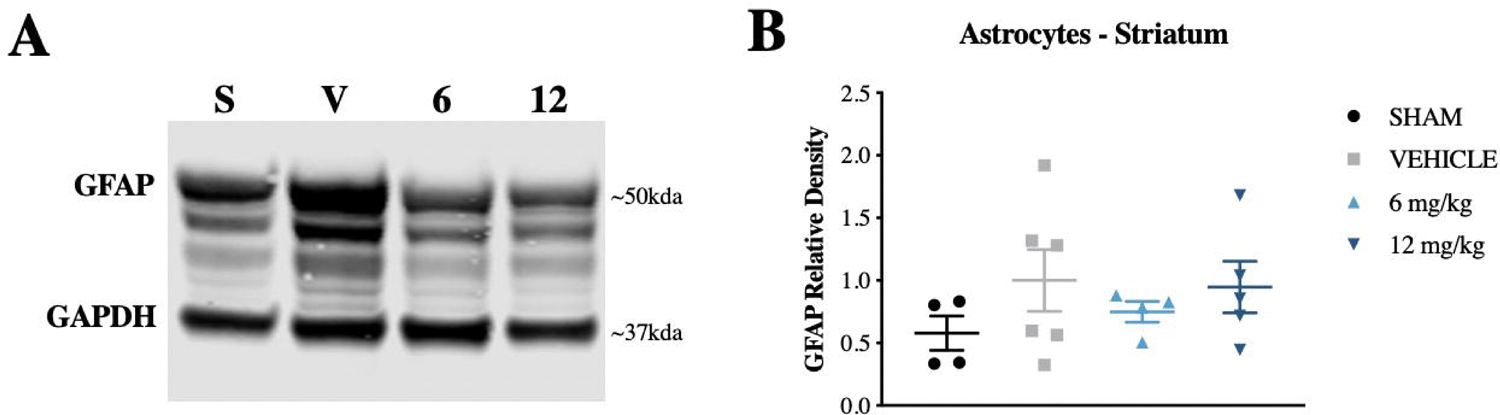
Astrocytic protein levels assessed by GFAP reactivity. (A) Western blot images of GFAP and GAPDH (housekeeper protein) markers. (B) Graph illustrates relative density of GFAP. Graph represents Mean± SEM (n=5-6 per group).

## 4. DISCUSSION

This was the first study to assess Fyn kinase inhibition as a therapeutic target for the treatment of PD. In order to trial this novel concept, the study aimed to investigate the efficacy of AZD0530 on a range of motor and non-motor outcomes and to assess the degree of neuroinflammation and dopaminergic integrity in an early stage model of PD. Results indicate animals receiving AZD0530 exhibited no significant changes in neuromuscular coordination, tremor or akinesia; however, they demonstrated improved volitional locomotion with 6 mg/kg and improved recognition memory and reduced depressive-like behaviour with both 6mg/kg and 12 mg/kg AZD0530 treatment. AZD0530 administration did not alter either striatal DA levels or inflammatory markers compared to vehicle-treatment, indicating alternate mechanisms or regionally specific changes may be responsible for improvements seen in behavioural outcomes.

In terms of motor outcomes, no impairments were observed in neuromuscular coordination or tremor in this early-stage model of PD, which corresponded to an average loss of only ∼30% of TH staining within the striatum. This is consistent with clinical findings in individuals with PD, where motor deficits manifest only when 70-80% of the DA terminals are destroyed and progressive DA neuronal loss has already occurred (Poston et al., 2016). Conversely, progressive motor impairments in both akinesia and volitional locomotion were observed with this model, appearing after 21 days, with improvements in volitional locomotion observed with AZD0530 treatment at day 33. These disparities in motor impairments and treatment outcomes may be attributed to subtle anatomical differences in striatal dopamine content affected in this model.

Striatal dopamine represents the main source of input for the basal ganglia and is therefore crucial for motor behaviour (Haber, 2014). The striatum itself contains 6 anatomical subdivisions, ventro/dorso-lateral, ventro/dorso-medial and ventro/dorso-central, each with connections to different cortical areas and, subsequently, these subdivisions are often associated with dominance in different motor functionalities (Kirik et al., 1998). For example, tremor is largely dependent on involvement of the ventro-lateral striatum (Collins-Praino et al., 2011), which predominantly remained intact in this early-stage model. In line with this, increases in tremor, as measured by TJMs, were not observed in this model. Similarly, neuromuscular coordination and sensorimotor learning, as measured by rotarod performance, have also been associated with ventro-lateral striatal lesions and also remained unaffected by intrastriatal 6-OHDA administration (Kirik et al., 1998).

Conversely, the AST, a measure of akinesia, is dependent on more widespread DA loss in the dorso-lateral, ventro-central and ventro-lateral striatum (Chang et al., 1999) and volitional locomotion in the OFT on dorso-lateral and dorso-medial striatal integrity (Barneoud et al., 2000).

An injury effect in the AST was seen at day 21 and in both the OFT and AST at day 33, suggesting that the dorsal DA loss seen in our model may be driving these disparate motor outcomes. Future work should investigate the ability of AZD0530 to treat motor impairments in a preclinical model involving DA loss to the ventro-lateral striatum, in order to more fully assess its benefits for treating motor symptoms of PD. Additionally, a longer time-point may be needed, given that previous literature utilising the intra-striatal 6-OHDA model has reported no changes in locomotion at 3 or 6 weeks post-lesion, but significantly less activity at 9 weeks (Branchi et al., 2008).

Given the significant impact of cognitive and neuropsychiatric impairments on quality of life for individuals with PD and their caregivers, we also assessed the ability of AZD0530 to target these non-motor symptoms. In this early-stage model, however, no significant impairments in learning, cognitive flexibility or visuo-spatial memory were observed on the Barnes Maze. This is in line with previous work in a comparable model, which also reported sparing of learning and visuo-spatial memory (Branchi et al., 2010; Branchi et al., 2008). Accordingly, the ability of AZD0530 to target such impairments could not be adequately assessed.

There were, however, significant improvements in recognition memory, as observed on the ORT. Compared to spatial or working memory, recognition memory is a relatively simplistic process (Broadbent et al., 2004); hence, it may be more sensitive to the subtle changes in DA signalling seen in this early-stage model. Recognition memory has also been shown to be significantly impaired in PD (Whittington et al., 2000) and connectivity studies have revealed an association with the progressive loss of normal network activity patterns in fronto-parietal connectivity (Segura et al., 2013). Whilst not well understood in the early 6-OHDA model, evidence suggests PD patients may exhibit impairments specifically in familiarity, rather than recollection (Davidson, 2006). Familiarity appears to be dependent on the medial temporal lobe (MTL) and extra-hippocampal structures (parahippocampal/perirhinal cortex) (Gabrieli et al., 1997). Additionally, a case study of a patient with a basal ganglia lesion demonstrated impairments in familiarity, but not recollection, implicating caudate damage may also underlie these impairments (Hay et al., 2002). Therefore, it may be beneficial for future studies to assess markers of inflammation and structural integrity in these areas.

This study also observed a significant decrease in depressive-like behaviour with both 6 mg/kg and 12 mg/kg treatment compared to vehicle. The early 6-OHDA model is known to recapitulate depressive-like behaviour, as was affirmed in our findings (Branchi et al., 2010). This is also in line with clinical data indicating depression may be a prominent non-motor symptom of prodromal PD, even preceding the motor symptoms of the disease (Jacob et al., 2010; Leentjens et al., 2003). In fact, depression is experienced by approximately 40% of PD patients early in disease progression (Tolosa et al., 2007). Anatomically, depression in early PD has been linked to dopamine-modulated fronto-striatal network dysfunction (Biundo et al., 2016). This network involves projections from the striatum to the orbitofrontal cortex, which is linked to regulation of emotion; consequently, loss of striatal DA may alter signalling throughout this network (Drevets, 2007). The reduction in depressive behaviour observed with AZD0530 may potentially represent conservation of DA function across this network or in other frontal regions.

Consistent with previous literature in this model (Thornton and Vink, 2012), increases in both microglial number and activation state were observed in all 6-OHDA-lesioned groups compared to shams in this study. Interestingly, however, there was no effect of AZD0530 treatment on either total number of microglia or microglial activation state. This is inconsistent with a previous study by Panicker and colleagues (2015), who established a decrease in microglial response in Fyn-/-6-OHDA lesioned mice, with associated nigral DA survival (Panicker et al., 2015). This may be dose-related, with a more substantial inhibition of Fyn kinase than that achieved with the dose regime in the current study potentially required to elicit a response. Similarly, a longer time course of treatment may be necessary in order to see improvements. In line with this, a previous study investigating AZD0530 treatment in the APP/PS1 mouse model of Alzheimer’s disease reported significant reductions in cortical microglial activation following 7 weeks of daily dosing at 5mg/kg (Kaufman et al. 2015). Alternatively, behavioural results observed in this study may be related to alternate mechanisms involving Fyn signalling. This would be consistent with previous reports from preclinical studies in Alzheimer’s disease, which suggest that improvements seen with AZD0530 treatment are due to the rescue of loss of synaptic density (Kaufman et al., 2015; Smith et al., 2018; Toyonaga et al., 2019).

It is important to note that the role of Fyn in the brain is diverse and widespread (Matrone et al., 2020). Therefore, other molecular mechanisms, particularly effects on specific neurotransmitter systems, may be involved in the beneficial effects of AZD0530 treatment on recognition memory and depressive-like behaviour in our experimental model of early-stage PD. For example, Fyn kinase plays a well-established role in the phosphorylation of N-methyl-d-aspartate (NMDA) glutamate receptors, in particular, the R2A and R2B subunits, and is subsequently implicated in upregulated NMDA function (Trepanier et al., 2012). Although crucial for many normal functions, in PD, hyperphosphorylation and the resulting overactivation of NMDA receptors is a known driver of glutamate excitotoxicity (Iovino et al., 2020; Truong et al., 2009). Additionally, increased levels of NR2B subunit phosphorylation have been observed not only in the striatum (Dunah et al., 2000; Oh et al., 1998), but also in the hippocampus of 6-OHDA lesioned rats (Rostas et al., 1996), which may help to account for the impairments in recognition memory seen in our 6-OHDA model. Whilst Fyn-mediated phosphorylation is vital for normal hippocampal LTP, there appears to be an optimal activity range and excessive activation or inhibition may impact performance (Parsons et al., 2007). In support of this, Fyn inhibition prevented phosphorylation of the NMDAR2B subunit in an animal model of AD, which was subsequently associated with rescuing of memory impairments (Kaufman et al., 2015). Thus, Fyn inhibition via AZD0530 may help to prevent pathological increases in NMDA R2A/R2B subunit phosphorylation, thereby reducing NMDA overactivity, subsequently rescuing recognition memory impairments, as observed in our study.

The effects of AZD0530 on depressive-like behaviour observed in the current study may also be attributable to actions on NMDA receptors, with several NMDA R2A/R2B antagonists exhibiting potential anti-depressant effects in PD studies, indicating that downregulation of NMDA phosphorylation may be beneficial for the treatment of depression (Vanle et al., 2018). Concurrently, Fyn has also been shown to interact with the metabotropic glutamate receptor subtype mGlu5, which is dense in both the striatum and limbic systems (Swanson et al., 2005). A variety of studies have suggested that mGlu5 is implicated in mood and anxiety symptoms, with antagonists of these receptors shown to be efficacious in preclinical models of depression (Belozertseva et al., 2007; Tallaksen-Greene et al., 1998; Tatarczynska et al., 2001; Testa et al., 1994). Recently, Fyn has been shown to form a complex with the mGlu5 receptors in striatal neurons in an animal model of depression, with Fyn kinase inhibition via PP2 significantly lowering increased expression of mGlu5 in this model (Mao and Wang, 2020). Thus, in the current study, anti-depressive effects of AZD0530 may be attributable to reduced glutamatergic signalling either via interactions with NMDAR2B subunits and/or post-synaptic mGlu5. Thus, future studies are warranted to investigate the specific mechanisms involved in interactions between Fyn and NMDA in anatomical regions associated with depression in PD.

Alternatively, another mechanism which may also account for the improvements in not only in depressive-like behaviour, but also recognition memory, seen in the current study is the 5-HT receptor 6 (5-HT_6_R). 5-HT_6_R has been implicated in the regulation of memory processes and, whilst its role in PD is not well understood, several studies have implicated 5-HT_6_R in learning and memory disorders, depression, and cognitive impairments associated with AD (Ivachtchenko et al., 2016; King et al., 2008). Given the symptomatic parallels and the observation that, unlike most 5-HT receptors, the 5-HT_6_ receptor is highly expressed in the striatum, it is possible the 5-HT_6_R may be implicated in PD. Paradoxically, 5-HT_6_R antagonists and agonists have both been shown to improve recognition memory in rodents using the ORT and depression using the FST (Kendall et al., 2011; King et al., 2008; King et al., 2004; Wesołowska and Nikiforuk, 2007). Recently, evidence suggests Fyn may play an important role in the signalling pathways of the 5-HT_6_R receptor, with Fyn and the 5-HT_6_R receptor colocalised and similarly distributed in the rat brain in the cortex, hippocampus and hypothalamus (although striatal co-localisation was not investigated) (Yun et al., 2007). Additionally, the expression of Fyn increased 5-HT_6_R expression, while 5-HT mediated activation of 5-HT_6_R increased Fyn phosphorylation (Yun et al., 2007). Thus, it is possible that Fyn inhibition reduced 5-HT_6_R phosphorylation, contributing to the rescue of recognition memory and reduction of depressive-like behaviour. This may represent a future area of investigation with regards to Fyn inhibition and its effects on 5-HT_6_R signalling.

Taken together, the findings of this study indicate Fyn kinase inhibition through AZD0530 may be beneficial for the treatment of non-motor symptoms of Parkinson’s Disease, currently a major area of clinical need. Interestingly, however, these effects do not appear to occur through anti-inflammatory mechanisms, although a more thorough investigation of neuroinflammatory response, including microglial phenotype and cytokine production, is necessary in order to fully rule this out. Instead, it may be that Fyn inhibition led to its beneficial effects via actions on glutamatergic or serotonergic signalling, thereby potentially rescuing the loss of synaptic density in related anatomical regions. Future studies are needed to more fully explore these effects, as this may lead to more targeted use of Fyn kinase inhibition as a treatment for non-motor symptoms of PD.

## Author contributions

BG performed experiments, conducted data analysis and drafted the manuscript. LC and BE assisted with data collection and analysis. SM and FC supervised the project and edited the manuscript. LCP designed the experiments, supervised the project, advised on data analysis and substantially edited the manuscript.

## Grant Support

This work was supported by a grant from the Commercial Acceleration Scheme from Adelaide Enterprise at the University of Adelaide.

## Acknowledgments

The authors wish to thank the AstraZeneca Open Innovation programme for generously providingAZD0530 for these experiments, as well as for technical input. They also wish to thank Dr Stephanie Plummer and Lisa Drew for their assistance with blinding during these experiments, as well as Dr Emma Thornton for input into study design in the early stages of this work.

## Notes

### Competing Interest Statement

The authors have declared no competing interest.

